# Personalized Prediction of Differences in fMRI Connectivity Estimates Across Preprocessing Pipelines

**DOI:** 10.64898/2026.11.04.763874

**Authors:** Yury Maximov, Aleksandr Volkov, Tamoghna Chattopadhyay, Paul M. Thompson, Ildar Gabitov

## Abstract

Longitudinal functional MRI of the human brain is valuable for tracking changes in functional connectivity (FC) associated with disease progression or treatment response. Even so, measured FC values depend on preprocessing, and nuisance regression and temporal filtering affect individuals differently. We examined whether we could use earlier scans to predict how preprocessing would affect a later scan. In 69 ADNI (Alzheimer’s Disease Neuroimaging Initiative) participants scanned three times, we measured how nuisance regression and filtering, alone and in sequence, changed the measured FC. A ridge regression model combined these responses with baseline FC from the third scan to predict the effect of reversing the processing order. Just four coefficients were shared across 79,800 connections. In leave-one-participant-out testing, mean squared error was 0.012, compared with 0.017 for predicting the same average response for everyone. It outperformed the best of seven alternative predictions chosen for each participant (0.013; normalized gain 0.069, 95% bootstrap interval 0.006–0.134), with lower error in 44 of 69 participants. About half the benefit of using each person’s earlier responses, rather than the average across participants, came from which connections changed. This advantage persisted after accounting for acquisition variables. The ADNI-trained model reduced mean error in ten Midnight Scan Club participants without refitting. These results support using personal preprocessing history to predict changes in connectivity estimates on newly acquired scans.

## 1 INTRODUCTION

Longitudinal fMRI studies use functional connectivity (FC) changes to assess disease progression and treatment response, including in neuromodulation trials for Alzheimer’s disease and depression [1, 2]. However, preprocessing steps affect FC measures differently across individuals. Accounting for this heterogeneity can improve prediction and reproducibility in task-based fMRI [3, 4]. Confound-regression benchmarks show that strategies vary in reducing motion effects while preserving network structure [5]. This raises a question: *does a person’s preprocessing response contain information that persists across visits?* If so, earlier scans may help predict a specified operation’s effect on FC estimates at a later visit. This could help distinguish preprocessing-related variation from longitudinal FC changes, without establishing whether either has a biological origin.

We consider a setting where the later scan has already been acquired and its functional connectivity (FC) estimated using an initial processing pipeline, which we call *baseline FC*. We then predict how those FC estimates would change with additional preprocessing steps. This differs from predicting future brain-connectivity changes [6] or identifying individuals by connectivity patterns across scans (connectome “fingerprinting”) [7]. We compare earlier preprocessing responses with baseline FC from the same scan and earlier visits as predictors. We test whether response magnitude alone explains individual differences or whether the connections affected provide additional information. This could help identify longitudinal FC findings robust to preprocessing choices, particularly when disease or treatment effects are small relative to preprocessing-related variation.

We tested this in 69 Alzheimer’s Disease Neuroimaging Initiative (ADNI) participants with three fMRI visits. Using the first two visits, we measured each person’s responses to nuisance regression and temporal filtering, then asked whether these responses could help predict the effect of reversing the processing order at the third visit. We fitted a ridge regression model with just four coefficients shared across all FC connections, and evaluated it by leaving out one participant at a time. To establish whether personal history was useful, we compared its predictions with seven alternatives, including those based on baseline FC and the average preprocessing response across participants. We examined whether the benefit came only from each person’s response magnitude or from the particular connections affected, and whether scanning protocol differences could explain the results. Finally, we applied the ADNI model without refitting to ten Midnight Scan Club (MSC) [8] participants to assess generalization to another dataset. Our aim was to predict preprocessing effects, rather than determine which pipeline is best for a specific objective, or whether these effects have a neural origin.

## 2 DATA AND METHODS

### 2.1 Participants and Cohort Selection

We analyzed ADNI resting-state fMRI data [9], initially consisting of 4,347 BOLD time series across 4,257 visits from 1,764 participants. Eligibility required a diagnosis recorded within 90 days of scanning, a suitable T1-weighted anatomical reference, reliable linkage to the original imaging data, and sufficient fMRI data. This left 3,810 visits from 1,698 participants, including 907 with at least two eligible visits. An index-visit T1-weighted scan from the same study and recorded sex information were required, leaving 903 participants. To balance the diagnostic groups, we matched each of the 90 eligible participants with dementia to one cognitively normal (CN) participant and one with mild cognitive impairment (MCI). Matching required an equal number of eligible visits, while accounting for acquisition and demographic characteristics. This gave us 270 participants with 732 visits.

Our prediction approach requires two earlier visits to establish each person’s preprocessing responses, and a third to test their reproducibility. Thus we focused on 69 participants (23 matched sets) with three eligible visits, retaining all 207 visits in chronological order. No missing responses were imputed or participants replaced. To evaluate prediction accuracy in new individuals, we held out one participant at a time, using the other 68 to fit the model.

We did not use diagnosis or measures of disease progression to predict preprocessing effects, instead testing what could be learned from each person’s imaging data alone. We compared predictions with several alternatives using the same 69 ADNI participants. We also tested the model in another dataset, using the first three sessions from ten Midnight Scan Club participants [8].

### 2.2 Preprocessing Responses and Prediction Target

We began with fMRI time series from 400 cortical regions defined by the Schaefer atlas [10], after regression of 24 head-motion parameters. Pearson correlations between regions gave 79,800 unique FC connections per visit. These estimates served as baseline FC (B) for assessing two preprocessing steps: anatomical CompCor (A), removing five nuisance-related components [11], and temporal high-pass filtering (F), implemented using a discrete cosine transform (DCT-II) with a 0.01 Hz cutoff. Because regression and filtering can be order-dependent [12], we tested five paths: baseline alone (BASE), CompCor alone (A), filtering alone (F), CompCor then filtering (AF), and filtering then CompCor (FA). To compare processing orders, we kept nuisance-regression components fixed rather than re-estimating them after filtering. In ADNI, we reconstructed these components from the difference between motion-regressed and joint motion-plus-CompCor residuals.

For each scan, we measured how much each preprocessing method changed FC relative to the baseline estimate from that same scan. We denote this change by Δ*O*_*it,e*_ where *O* identifies the preprocessing method, *i* the participant, *t* the visit, and *e* the FC connection. We used the first two visits to measure each person’s average response to nuisance regression (A), filtering (F), and their sequential application (AF). These three measurements, together with baseline FC from the third visit, provided four inputs for predicting each connection:

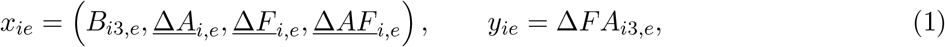

The bars show averages across visits 1 and 2, used to predict the FC change at visit 3 under the reversed preprocessing order (FA). Predicting the effect of a different processing sequence, rather than reproducing an earlier measured response, tests whether individual preprocessing sensitivity transfers across processing choices. Fig. 1 shows the prediction inputs and the response withheld for testing.

**Figure 1.**
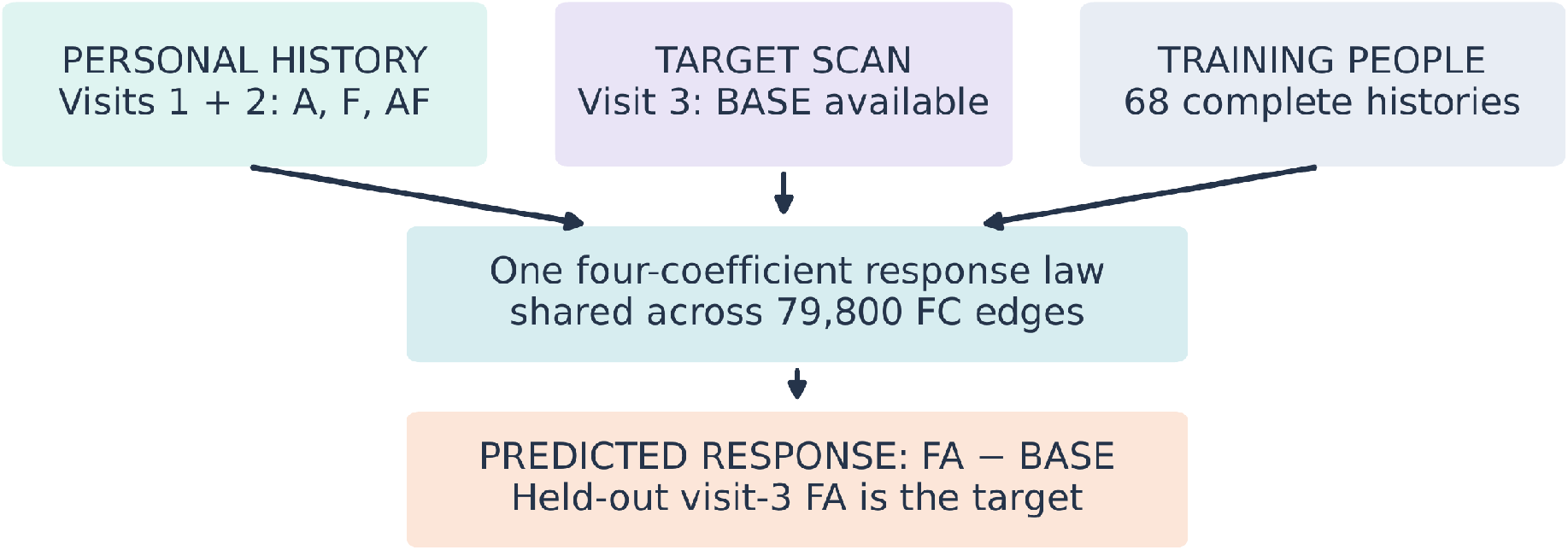
Prediction model. Preprocessing responses from visits 1 and 2 and baseline functional connectivity from visit 3 predict how reversing the order of nuisance regression and temporal filtering affects connectivity at visit 3. The actual third-visit response is withheld during prediction and used to evaluate accuracy.

### 2.3 Prediction Model and Controls

We used ridge regression to predict the third-visit preprocessing effect from baseline FC at that visit and the participant’s earlier responses. To assess generalization to new individuals, we used leave-oneparticipant-out cross-validation, fitting the model on the other 68 participants. Each of the 79,800 FC connections contributed a training observation, but the model used only 4 coefficients shared across all connections, estimated as

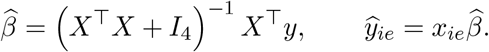

Here, *X* contains the four predictors from Eq. (1), and y the observed preprocessing responses. We used a ridge penalty of 1 without an intercept and measured mean squared error (MSE) across FC connections for each held-out participant, in squared-correlation units.

We next asked whether personal preprocessing history improved prediction over simpler alternatives. We compared the model with seven alternatives based on the average response across other participants; earlier regression and filtering responses, separately or in sequence; baseline FC from earlier or current scans; and summaries of scan reliability or preprocessing operations. Coefficients for these alternatives were fitted using training participants only. For a more demanding comparison, we compared our model with the lowest-error alternative for each participant. Selected using the observed response, this best-case benchmark is stricter than a practical comparison. We quantified improvement using mean losses *L*_env_, *L*_personal_ and *L*_univ_; the contrast is

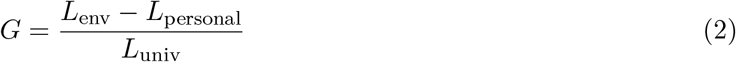

where *L*_env_ is the best-case benchmark’s mean error, *L*_personal_ is our model’s mean error, and *L*_univ_ is the error from predicting the population-average response for everyone. We calculated 95% bootstrap intervals from 20,000 resamples of participants’ prediction errors; the population-average error remained fixed, and models were not refitted.

First, we tested whether repeated baseline FC or acquisition-protocol differences could explain the results, selecting acquisition-related predictors using training data only. We compared personal preprocessing history with the across-participant average. To distinguish overall response strength from connection-specific differences, we tested predictions that scaled the average response pattern to each participant, keeping fitted coefficients unchanged. Finally, we tested whether agreement between visits 1 and 2 could identify participants most likely to benefit from personalized prediction. To test the model on another dataset, we fitted its four coefficients using all 69 ADNI participants and applied them without refitting to ten MSC participants [8].

## 3 RESULTS

### 3.1 Predicting Preprocessing Effects from Earlier Visits

We first tested whether a person’s FC responses to preprocessing at two earlier visits improved prediction at the third visit. The 4-coefficient model achieved a mean squared error (MSE) of 0.012 with each of the 69 participants held out in turn. Predicting the same average preprocessing response for everyone gave an MSE of 0.017. Among seven alternatives, the most accurate on average used baseline FC from the third scan (MSE = 0.015). For a more demanding comparison, we selected the lowest-error alternative for each participant after seeing their actual response. Even against this unusually favorable benchmark (MSE = 0.013), our model performed better, with lower error in 44 of 69 participants (Fig. 2). The improvement, normalized by the error of predicting the population-average response (Eq. (2)), was 0.069 (95% bootstrap interval, 0.006–0.134).

**Figure 2.**
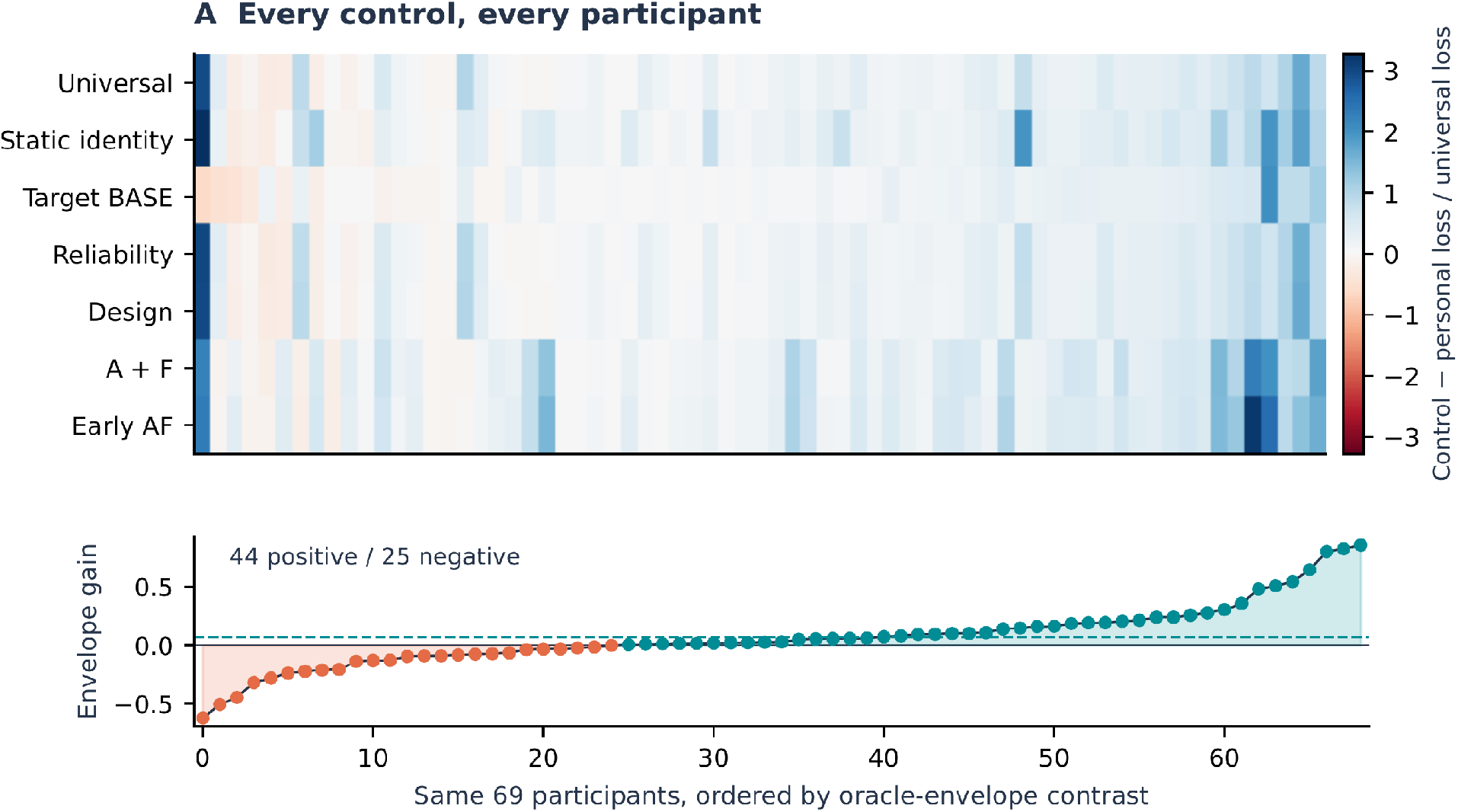
Prediction accuracy across 69 ADNI participants. (A) Personalized model versus seven alternatives. Columns show participants; rows show alternatives. Blue indicates lower error for the personalized model; red for the alternative. Differences are normalized by the error of predicting the average response across participants. (B) Improvement over each participant’s best of seven alternatives. Positive values favor the personalized model (44 of 69 participants). The dashed line shows mean improvement.

We tested whether scanning protocol differences could explain this improvement. After considering acquisition variables, the model still outperformed alternatives, with a normalized gain of 0.064 (95% interval, 0.001–0.128) and lower error in 41 participants. Acquisition information alone did not improve prediction over the population average (normalized gain −0.077, interval −0.277 to 0.102), nor did adding it to the original model (−0.011, interval −0.049 to 0.028). Thus, the acquisition variables examined did not explain the benefit of earlier preprocessing responses.

### 3.2 Contributions of Earlier Preprocessing Responses

We examined why earlier preprocessing responses improved prediction. Possibly, individuals may have relatively stable baseline FC patterns across visits, regardless of preprocessing. However, predictions based on repeated baseline FC had an MSE of 0.013, and including each person’s earlier preprocessing responses reduced this error by 4.3%, improving prediction in 40 of 69 participants.

Preprocessing may also affect everyone similarly, making the average response across participants sufficient. Replacing individual preprocessing histories with the training-participant average yielded an MSE of 0.013, higher than the personalized model. A 14-feature regression including quadratic terms also achieved an MSE of 0.012 (rounded), but the four-coefficient model had the lowest mean error.

To understand what was gained from using each person’s own history, we kept the fitted coefficients unchanged and varied only earlier-visit information. Let *b*_*i*_, *q*_*i*_, and *m*_*i*_ be the prediction based on the third scan’s baseline FC, the contribution from that person’s earlier preprocessing responses, and the average contribution from the training participants, respectively. Let *r*_*i*_ = *y*_*i*_ − *b*_*i*_ − *m*_*i*_, *δ*_*i*_ = *q*_*i*_ − *m*_*i*_ be the part of the observed response not explained by baseline FC and the average history, and the correction from personal history, respectively. For *D* = 79, 800 edges, the reduction in prediction error is:

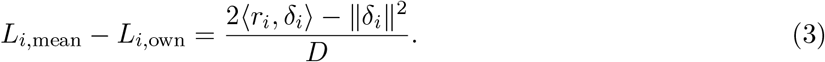

The first term measures whether the personal correction moves the prediction toward the observed response; the second penalizes corrections that are too large. Averaged across participants, these terms contributed 0.002 and 0.001, giving a net reduction in error of 0.001. Using personal rather than average preprocessing history lowered MSE by 6.7%. A further comparison using response histories from other participants gave an MSE of 0.014; 11.8% higher than that of the personal model. Variation between these other participants contributed ∼ 0.001 to the error.

Finally, we determined whether individuals differed mainly in how strongly they responded to preprocessing, or also in which FC connections were affected. We tested this by allowing the average response pattern to be scaled up, down, or reversed for each participant. This gave an MSE of 0.013, while retaining the person’s actual response pattern lowered error by a further 3.5%. The individual spatial pattern improved prediction in 41 participants. Overall, differences in response magnitude accounted for 49.7% of the improvement over the average pattern; differences in which connections were affected accounted for 50.3% (**Fig. 3**). The analysis retained all 14 participants whose fitted scaling factors were negative.

**Figure 3.**
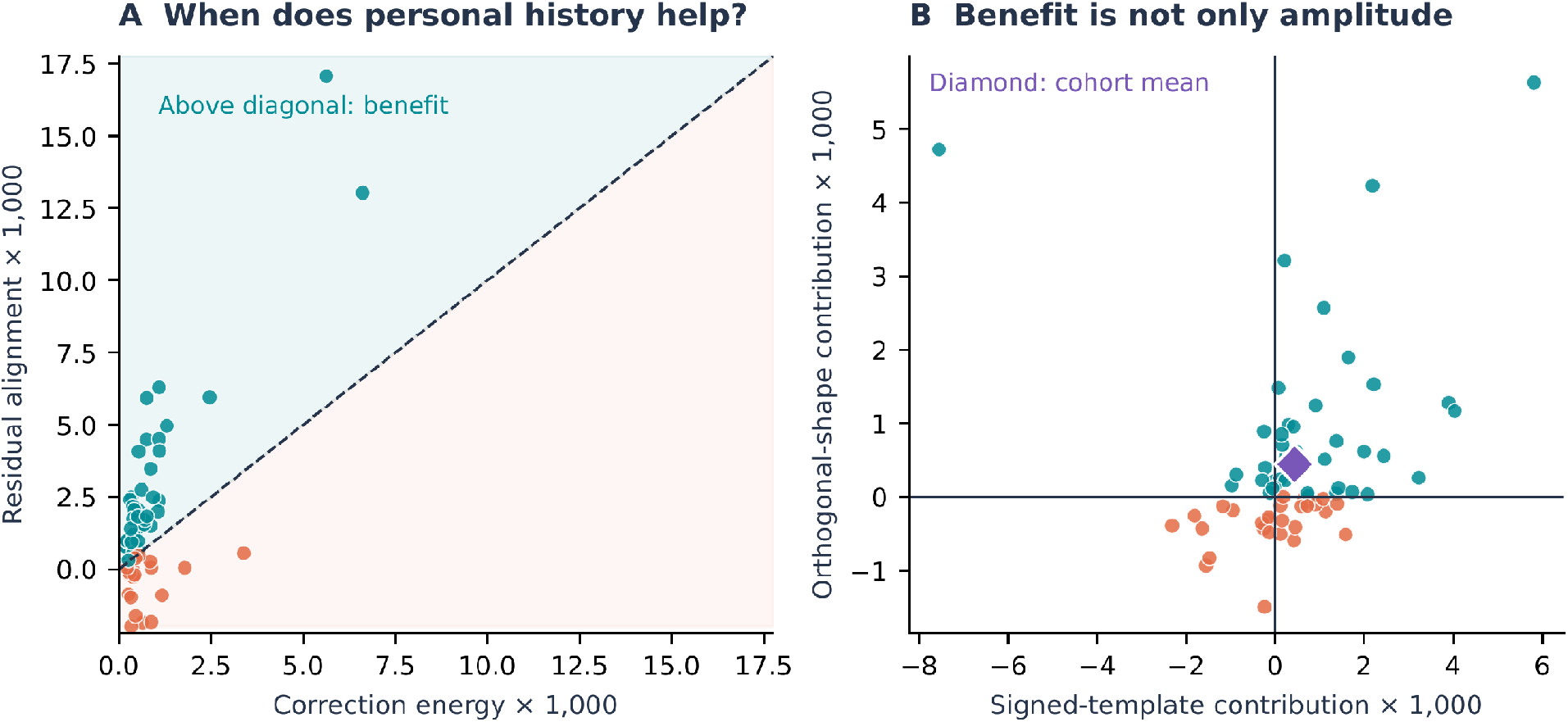
How individual preprocessing responses improve prediction. (A) For each participant, we compare personal-history prediction accuracy with FC correction magnitude. Points above the diagonal favor personal history over the across-participant average. (B) We separate this improvement into differences in overall response strength (horizontal axis) and affected FC connections (vertical axis). The diamond marks the participant average. Values are in squared-correlation units (×1,000).

### 3.3 Cross-Visit Agreement and Cross-Dataset Testing

Agreement across visits was positively correlated with improvement over repeated baseline FC (Spearman correlation = 0.306) and average preprocessing history (0.388). We tested whether this information could improve prediction by choosing between the personalized model and repeated baseline FC based on agreement between the first two visits. This approach selected personalized predictions for 40 participants and repeated baseline FC for 29, improving accuracy in only 12 and worsening it in 17. Overall MSE was 0.013, 2.77% higher than using the personalized model for everyone. Thus, agreement was informative but did not improve model selection.

To assess cross-dataset generalization, we fitted the four coefficients using all 69 ADNI participants and applied them without refitting to ten MSC participants. The personalized model achieved an MSE of 0.004, compared with 0.0046 for repeated baseline FC, 0.0048 for average ADNI preprocessing history, and 0.0053 for the average ADNI response at the third visit. Error reductions were 5.3%, 9.2%, and 18.7%, respectively. Personal history improved prediction over repeated baseline FC in 6 of 10 participants and over average history in 8 (**Fig. 4**).

**Figure 4.**
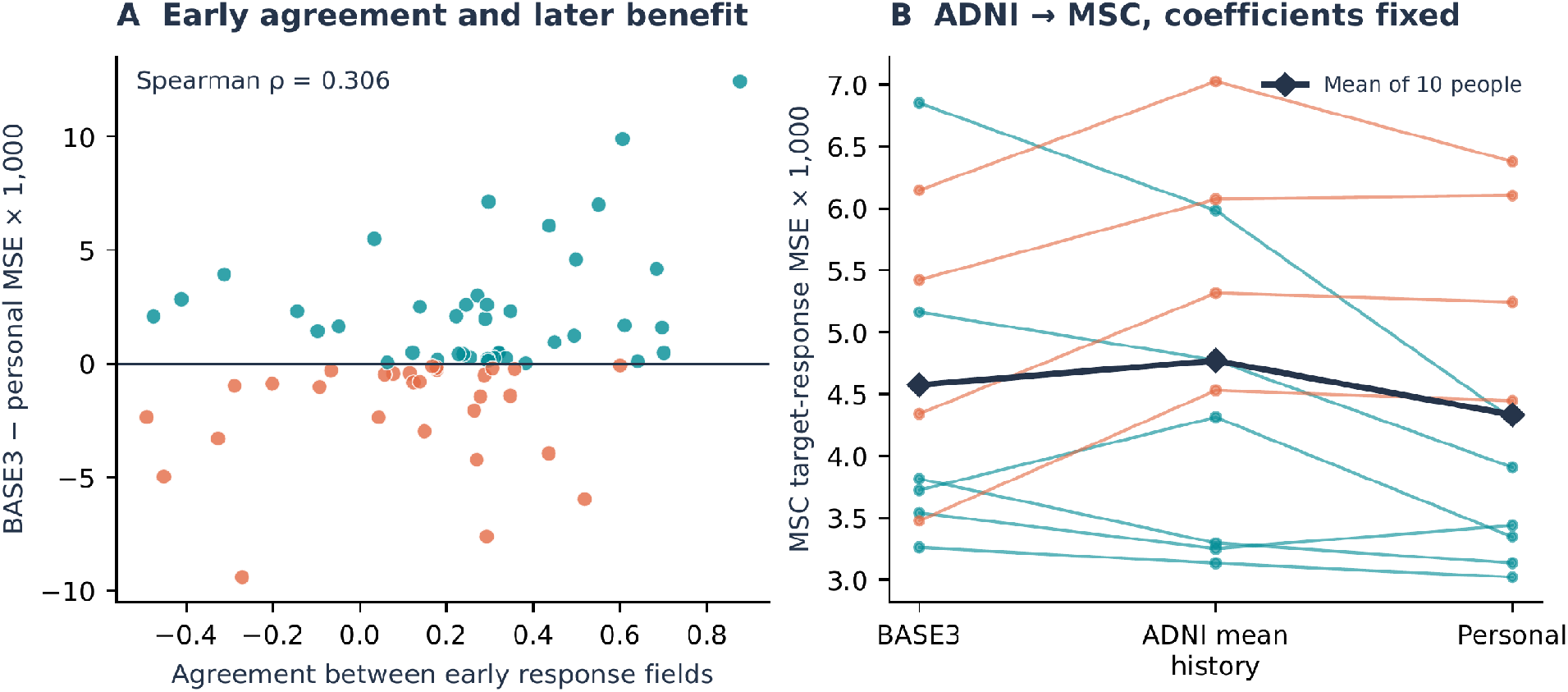
Cross-visit consistency and cross-dataset prediction. (A) Greater agreement between preprocessing responses at visits 1 and 2 was associated with greater benefits from personalized prediction at visit 3. (B) Prediction errors for ten MSC participants, using ADNI-trained model coefficients without refitting. Lines connect each participant’s errors for repeated baseline FC, average ADNI preprocessing history, and personal preprocessing history. Diamonds show the mean errors across participants.

Finally, we tested whether baseline FC patterns alone could identify participants across visits. Earlier baseline patterns correctly identified 59 of 69 third visits. This separate analysis was not used to select or evaluate the preprocessing prediction model.

## 4 DISCUSSION & LIMITATIONS

We found that a person’s earlier responses to fMRI preprocessing can help predict how a different processing sequence affects their connectivity estimates at a later visit. A model with four shared coefficients outperformed predictions based on average preprocessing responses or repeated baseline FC. It outperformed, on average, the best of seven alternatives selected separately for each participant. About half the gain from personal history reflected which connections were affected rather than response magnitude. This suggests individual-specific patterns of preprocessing sensitivity persist across visits.

These findings matter for longitudinal fMRI studies, where small FC changes may be interpreted as evidence of disease progression or treatment response. If preprocessing affects individuals reproducibly, earlier scans may help assess findings’ dependence on processing choices. We extend prior work optimizing preprocessing pipelines [4] and comparing their effects on FC [13]; we show that individual responses are predictable across visits even when preprocessing order changes.

Several limitations remain. Reproducible preprocessing responses may reflect systematic individual differences in physiological signals or head motion rather than stable neural properties [14, 15]. Our results do not establish the biological origin of the responses or indicate which preprocessing method is preferable. Finally, bootstrap intervals reflect uncertainty in observed prediction errors, excluding model-refitting uncertainty.

Overall, preprocessing sensitivity could be predicted from an individual’s earlier scans using a relatively simple model. Improvements persisted after accounting for acquisition differences. Larger, independent longitudinal datasets are needed to assess generalizability and whether accounting for preprocessing sensitivity improves the reliability of disease progression and treatment response studies.

## 5 CODE AND DATA AVAILABILITY

Code for the prediction model, the comparison methods, and the example applying the ADNI-trained model to MSC is available on GitHub: https://github.com/yury-rd/fmri-transfer

## 6 COMPLIANCE WITH ETHICAL STANDARDS

We analyzed previously collected, publicly available ADNI and MSC data, without recruiting participants or acquiring new scans. For this analysis of public, de-identified anonymized data, no additional institutional review board approval was required. Ethical procedures for the MSC study are described in the original publication [8] and the representative MSC dataset webpage https://openneuro.org/datasets/ds000224/versions/1.0.4.

## Acknowledgments

Data used in preparation of this article were obtained from the Alzheimer’s Disease Neuroimaging Initiative (ADNI) database (adni.loni.usc.edu). As such, the investigators within the ADNI contributed to the design and implementation of ADNI and/or provided data but did not participate in the analysis or writing of this report. A complete listing of ADNI investigators can be found in the ADNI investigator acknowledgment list.

The ADNI imaging and clinical data came from the Alzheimer’s Disease Neuroimaging Initiative database. ADNI began in 2003 as a public-private partnership led by Michael W. Weiner, MD. It brings together longitudinal imaging, biomarkers, and clinical and neuropsychological assessments to study progression from mild cognitive impairment to Alzheimer’s disease.

The National Institute on Aging funds ADNI data collection and sharing through National Institutes of Health grant U19 AG024904 to the Northern California Institute for Research and Education. ADNI has also received support from the National Institute of Biomedical Imaging and Bioengineering, the Canadian Institutes of Health Research, and private contributors through the Foundation for the National Institutes of Health. These contributors include AbbVie, Alzheimer’s Association, Alzheimer’s Drug Discovery Foundation, Araclon Biotech, BioClinica, Inc., Biogen, Bristol-Myers Squibb Company, CereSpir, Inc., Cogstate, Eisai Inc., Elan Pharmaceuticals, Inc., Eli Lilly and Company, EuroImmun, F. Hoffmann-La Roche Ltd and its affiliated company Genentech, Inc., Fujirebio, GE Healthcare, IXICO Ltd., Janssen Alzheimer Immunotherapy Research & Development, LLC., Johnson & Johnson Pharmaceutical Research & Development LLC., Lumosity, Lundbeck, Merck & Co., Inc., Meso Scale Diagnostics, LLC., NeuroRx Research, Neurotrack Technologies, Novartis Pharmaceuticals Corporation, Pfizer Inc., Piramal Imaging, Servier, Takeda Pharmaceutical Company, and Transition Therapeutics.

## References

[1] Y. H. Jung, H. Jang, S. Park, H. J. Kim, S. W. Seo, G. B. Kim, Y.-M. Shon, S. Kim, and D. L. Na, “Effectiveness of Personalized Hippocampal Network-Targeted Stimulation in Alzheimer Disease: A Randomized Clinical Trial,” JAMA Network Open, vol. 7, no. 5, Art. no. e249220, 2024. doi: 10.1001/jamanetworkopen.2024.9220.

[2] J.-M. Batail, X. Xiao, A. Azeez, C. Tischler, I. H. Kratter, J. H. Bishop, M. Saggar, and N. R. Williams, “Network Effects of Stanford Neuromodulation Therapy (SNT) in Treatment-Resistant Major Depressive Disorder: A Randomized, Controlled Trial,” Translational Psychiatry, vol. 13, no. 1, Art. no. 240, 2023. doi: 10.1038/s41398-023-02537-9.

[3] N. W. Churchill, G. Yourganov, A. Oder, F. Tam, S. J. Graham, and S. C. Strother, “Optimizing Preprocessing and Analysis Pipelines for Single-Subject fMRI: 2. Interactions with ICA, PCA, Task Contrast and InterSubject Heterogeneity,” PLOS ONE, vol. 7, no. 2, Art. no. e31147, 2012. doi: 10.1371/journal.pone.0031147.

[4] N. W. Churchill, R. Spring, B. Afshin-Pour, F. Dong, and S. C. Strother, “An Automated, Adaptive Framework for Optimizing Preprocessing Pipelines in Task-Based Functional MRI,” PLOS ONE, vol. 10, no. 7, Art. no. e0131520, 2015. doi: 10.1371/journal.pone.0131520.

[5] R. Ciric, D. H. Wolf, J. D. Power, D. R. Roalf, G. L. Baum, K. Ruparel, R. T. Shinohara, M. A. Elliott, S. B. Eickhoff, C. Davatzikos, R. C. Gur, R. E. Gur, D. S. Bassett, and T. D. Satterthwaite, “Benchmarking of Participant-Level Confound Regression Strategies for the Control of Motion Artifact in Studies of Functional Connectivity,” NeuroImage, vol. 154, pp. 174–187, 2017. doi: 10.1016/j.neuroimage.2017.03.020.

[6] W. Xia, X. Zhang, D. Hu, J. Cheng, Z. Wu, L. Wang, W. Lin, and G. Li, “Individualized Trajectory Prediction of Early Developing Functional Connectivity,” in 2025 IEEE 22nd International Symposium on Biomedical Imaging (ISBI), pp. 1–5, 2025. doi: 10.1109/ISBI60581.2025.10980810.

[7] E. S. Finn, X. Shen, D. Scheinost, M. D. Rosenberg, J. Huang, M. M. Chun, X. Papademetris, and R. T. Constable, “Functional Connectome Fingerprinting: Identifying Individuals Using Patterns of Brain Connectivity,” Nature Neuroscience, vol. 18, no. 11, pp. 1664–1671, 2015. doi: 10.1038/nn.4135.

[8] E. M. Gordon, T. O. Laumann, A. W. Gilmore, D. J. Newbold, D. J. Greene, J. J. Berg, M. Ortega, C. Hoyt-Drazen, C. Gratton, H. Sun, J. M. Hampton, R. S. Coalson, A. L. Nguyen, K. B. McDermott, J. S. Shimony, A. Z. Snyder, B. L. Schlaggar, S. E. Petersen, S. M. Nelson, and N. U. F. Dosenbach, “Precision Functional Mapping of Individual Human Brains,” Neuron, vol. 95, no. 4, pp. 791–807.e7, 2017. doi: 10.1016/j.neuron.2017.07.011.

[9] C. R. Jack, Jr., M. A. Bernstein, N. C. Fox, P. Thompson, G. Alexander, D. Harvey, B. Borowski, P. J. Britson, J. L. Whitwell, C. Ward, A. M. Dale, J. P. Felmlee, J. L. Gunter, D. L. G. Hill, R. Killiany, N. Schuff, S. Fox-Bosetti, C. Lin, C. Studholme, C. S. DeCarli, G. Krueger, H. A. Ward, G. J. Metzger, K. T. Scott, R. Mallozzi, D. Blezek, J. Levy, J. P. Debbins, A. S. Fleisher, M. Albert, R. Green, G. Bartzokis, G. Glover, J. Mugler, and M. W. Weiner, “The Alzheimer’s Disease Neuroimaging Initiative (ADNI): MRI Methods,” Journal of Magnetic Resonance Imaging, vol. 27, no. 4, pp. 685–691, 2008. doi: 10.1002/jmri.21049.

[10] A. Schaefer, R. Kong, E. M. Gordon, T. O. Laumann, X.-N. Zuo, A. J. Holmes, S. B. Eickhoff, and B. T. T. Yeo, “Local-Global Parcellation of the Human Cerebral Cortex from Intrinsic Functional Connectivity MRI,” Cerebral Cortex, vol. 28, no. 9, pp. 3095–3114, 2018. doi: 10.1093/cercor/bhx179.

[11] Y. Behzadi, K. Restom, J. Liau, and T. T. Liu, “A Component Based Noise Correction Method (CompCor) for BOLD and Perfusion Based fMRI,” NeuroImage, vol. 37, no. 1, pp. 90–101, 2007. doi: 10.1016/j.neuroimage.2007.04.042.

[12] M. A. Lindquist, S. Geuter, T. D. Wager, and B. S. Caffo, “Modular Preprocessing Pipelines Can Reintroduce Artifacts into fMRI Data,” Human Brain Mapping, vol. 40, no. 8, pp. 2358–2376, 2019. doi: 10.1002/hbm.24528.

[13] A. I. Luppi, H. M. Gellersen, Z.-Q. Liu, A. R. D. Peattie, A. E. Manktelow, R. Adapa, A. M. Owen, L. Naci, D. K. Menon, S. I. Dimitriadis, and E. A. Stamatakis, “Systematic Evaluation of fMRI Data-Processing Pipelines for Consistent Functional Connectomics,” Nature Communications, vol. 15, no. 1, Art. no. 4745, 2024. doi: 10.1038/s41467-024-48781-5.

[14] S. Noble, D. Scheinost, and R. T. Constable, “A Decade of Test-Retest Reliability of Functional Connectivity: A Systematic Review and Meta-Analysis,” NeuroImage, vol. 203, Art. no. 116157, 2019. doi: 10.1016/j.neuroimage.2019.116157.

[15] A. Xifra-Porxas, M. Kassinopoulos, and G. D. Mitsis, “Physiological and Motion Signatures in Static and Time-Varying Functional Connectivity and Their Subject Identifiability,” eLife, vol. 10, Art. no. e62324, 2021. doi: 10.7554/eLife.62324.

